# Find, Label, Annotate Genomes: FLAG, a fully automated tool for structural and functional gene annotation

**DOI:** 10.1101/2023.07.14.548907

**Authors:** William Troy, Joana Damas, Alexander J. Titus, Brandi L. Cantarel

## Abstract

Recent advances in long-read sequencing technologies and the efforts of projects aimed at increasing the universe of sequenced reference genomes have led to a growth in the number of whole genomes sequenced for non-model organisms. Still, 81% of the over 36,000 unique publicly available eukaryotic genomes in the NCBI database lack gene structure annotations (1). While there are many open-source tools available for each step in the annotation process, many of these tools are designed for chromosomal assemblies with available transcript data from the same or very closely related organisms. Here we present “Find, Label, Annotate Genomes” (FLAG), a fully automated genome annotation workflow. FLAG (i) works on any computing environment, (ii) runs automatically without initial training data, (iii) generates structural and functional annotations, (iv) performs accurately with fragmented genomes, (v) does not require species-specific extrinsic evidence (transcript sequences) and (vi) includes quality control steps to evaluate annotation completeness. We compared the gene annotations generated by FLAG and publicly available annotations from 12 eukaryotic organisms, including 1 plant, 8 living animals, and 1 extinct animal. In fragmented genomes, FLAG annotations provided an average of 18% increase in complete BUSCO scores and 15x lower error rate for the predicted number of protein-coding genes when compared to published BRAKER2 annotations. With FLAG-Refiner further improved results by decreasing the error rate to 17x lower when compared to published BRAKER2 annotations. In high-quality model organisms, FLAG demonstrates comparable results to those of the NCBI EGAP pipeline, underscoring its robustness and accuracy in gene prediction across diverse taxa and assembly qualities.

## Introduction

Due to the highly repetitive and complex nature of eukaryotic genomes, accurately finding genes is an active area of research. The last five years have seen great advances in sequencing technologies that led to the availability of more accurate and complete genome assemblies.

Structural eukaryotic genome annotation can involve gathering transcript sequencing data for the species of interest, comparison of genes from closely related species, and iterating genome gene structure annotation following training gene models based on “high confidence” gene models (2– 7). Genome annotation quality is highly dependent on genome assembly accuracy and completeness (4, 8). Genome annotation accuracy is decreased when (i) sequences are missing; (ii) assemblies are highly fragmented, which can fragment genes on multiple contigs, and (iii) contigs are misassembled. Typical crustacean and tree genomes that are of low quality or are highly gapped have over 100k extra protein-coding genes when compared to complete annotations of related species (9–11), indicating a large error rate in their annotations.

End-to-end workflows such as BRAKER2 (4, 7) and MAKER (5, 12) can take supporting evidence to train gene-finding tools such as Gnomon, Augustus, GeneMark-ES, or SNAP (4, 5, 7, 13) and use those models to produce gene predictions. BRAKER2 is the genome annotation workflow used by Ensembl Fast Track and has been shown to be more accurate than previously published eukaryotic annotation workflows (4). These methods require large computational resources and have long runtimes, usually requiring >24h for training only, tend to be less accurate for highly fragmented genomes, and require some computational expertise to install and run. Finally, while the NCBI EGAP pipeline (13) is considered the gold standard of gene annotation and is regularly used for the benchmarking of newly developed tools (4, 14, 15).

However, as of writing this paper, it is not fully publicly available. On top of this, the NCBI only has the resources to annotate ∼200 genomes per year, to which they require transcript data from multiple tissues and are generally only on genome assemblies of the highest quality (1, 16). All of this together creates a clear gap in computational genomics where there is a need for a fast, highly accurate, genome annotation pipeline that does not require species-specific evidence and can annotate not just high-quality genomes but also those of lesser quality.

To alleviate this problem we developed “Find, Label, Annotate Genomes” (FLAG) as an end-to-end automated workflow to accurately annotate eukaryotic genomes even when extrinsic or homology-based data is nonexistent. FLAG can accept species-specific extrinsic evidence and homology-based training data but can also create highly complete and accurate annotations when this data is unavailable. FLAG allows the user to combine results from homology-based alignments and multiple gene finding methods including Liftoff (17), Augustus (2), and Helixer(8). We combine the results of all gene predictors using a novel combine and filter algorithm, functionally annotate, and, when possible, assign a gene symbol to the resulting gene models using Entap (18) or Eggnog (19). Finally, these annotations are evaluated with FLAG-Refiner, a novel tool that we developed to determine the confidence of gene predictions. FLAG-Refiner compares gene predictions against a large database curated from all public annotations in the RefSeq and Ensembl databases as well as all protein- and transcript-to-genome alignments produced in a given run of FLAG. All FLAG results are also paired with AGAT (20) and BUSCO (21, 22) reports for further evaluation of genome annotation quality. AGAT is used to determine general annotation statistics such as the number of annotations of various types (e.g., protein-coding, mRNA, tRNA) along with their minimum, maximum, and average annotation lengths. BUSCO can then be used to identify the presence of highly conserved genes within a given lineage allowing us to infer the approximate accuracy of the overall genome annotation.

This evaluation of the accuracy of conserved genes is particularly useful when AGAT metrics align with expected standards as it can then roughly be extrapolated to determine overall gene annotation accuracy.

Furthermore, in our comparative analysis with the EGAP pipeline from NCBI, we leveraged FLAG-Refiner’s capabilities to determine the likeliness of gene annotations by incorporating external evidence. This approach proved instrumental in situations where the predicted gene counts diverged from those provided by EGAP, offering an added layer of validation and support to our findings. This dual approach of employing AGAT for detailed metrics and BUSCO for conservation-based validation, supplemented by FLAG-Refiner’s evidence integration, provides a comprehensive framework for assessing the efficacy of genome annotation workflows, highlighting the robustness and accuracy of FLAG in annotating genomes across diverse qualities.

## Materials and Methods

Non-model species assemblies annotated in this study along with their BRAKER2 annotations include the dingy skipper butterfly (*Erynnis tages* (23)), scarce swallowtail butterfly (*Iphiclides podalirius* (24)), iron walnut tree (*Juglans sigillata* (10)), golden perch (*Macquaria ambigua* (25)), Japanese white-eye (*Zosterops japonicus* (26)), Kordofan giraffe (*Giraffa camelopardalis antiquorum* (27)), and little-bush moa (*Anomalopteryx didiformis* (28)). Model RefSeq genome assemblies annotated along with their corresponding NCBI EGAP annotations can be found in the RefSeq database with the following accession numbers: Anna’s hummingbird (*Calypte anna*; GCF_003957555.1), chicken (*Gallus gallus;* GCF_016699485.2), and zebra finch (*Taeniopygia guttata;* GCF_003957565.2). Accession information on genome assemblies and annotations used as references, tool speed tests, and evidence for annotations can be found in Supplementary Table 1.

### Input Data Types

Genome annotation requires two main types of input: a target genome sequence and external evidence. For external evidence, FLAG accepts any combination of RNAseq, transcript and protein sequences from the species of interest and a closely related species, as well as the genome sequence and corresponding gene annotations of a closely related species. Genome assemblies (FastA) and annotations (GFF or GTF) from closely related species should be of good quality and completeness, and from species with 95% or higher genome sequence identity.

While it is best to supply transcript or protein data from the organism of interest, if that data is not available, users can use publicly available data for a closely related species, usually available from RefSeq (29), UniProt (30), and UniRef50 (31). Users of the Form Bio platform will be able to select from a set of curated clade-specific databases such as mammalian, avian, sauropsids, insect, and fish, among others.

### Annotation Methods

Based on user selection and input data, FLAG will run different combinations of tools to determine genes (i) that are homologous to sequencing data from the target or a closely related species, (ii) share excess similarity to proteins in a protein reference database and/or (iii) are predicted using ab-initio gene finding methods (Figure 1). FLAG has three main sub-workflows: (i) the RNA workflow, which can create a transcript assembly and align transcripts to the target genome, (ii) the protein workflow, which identifies putative genes by homology with known protein sequences, and (iii) the gene finding workflow that uses evidence from the RNA and protein workflows to inform on ab-initio gene predictions, synthesize all evidence into gene models, and annotate genes with orthologous genes name (Figure 2).

**Figure 1.**
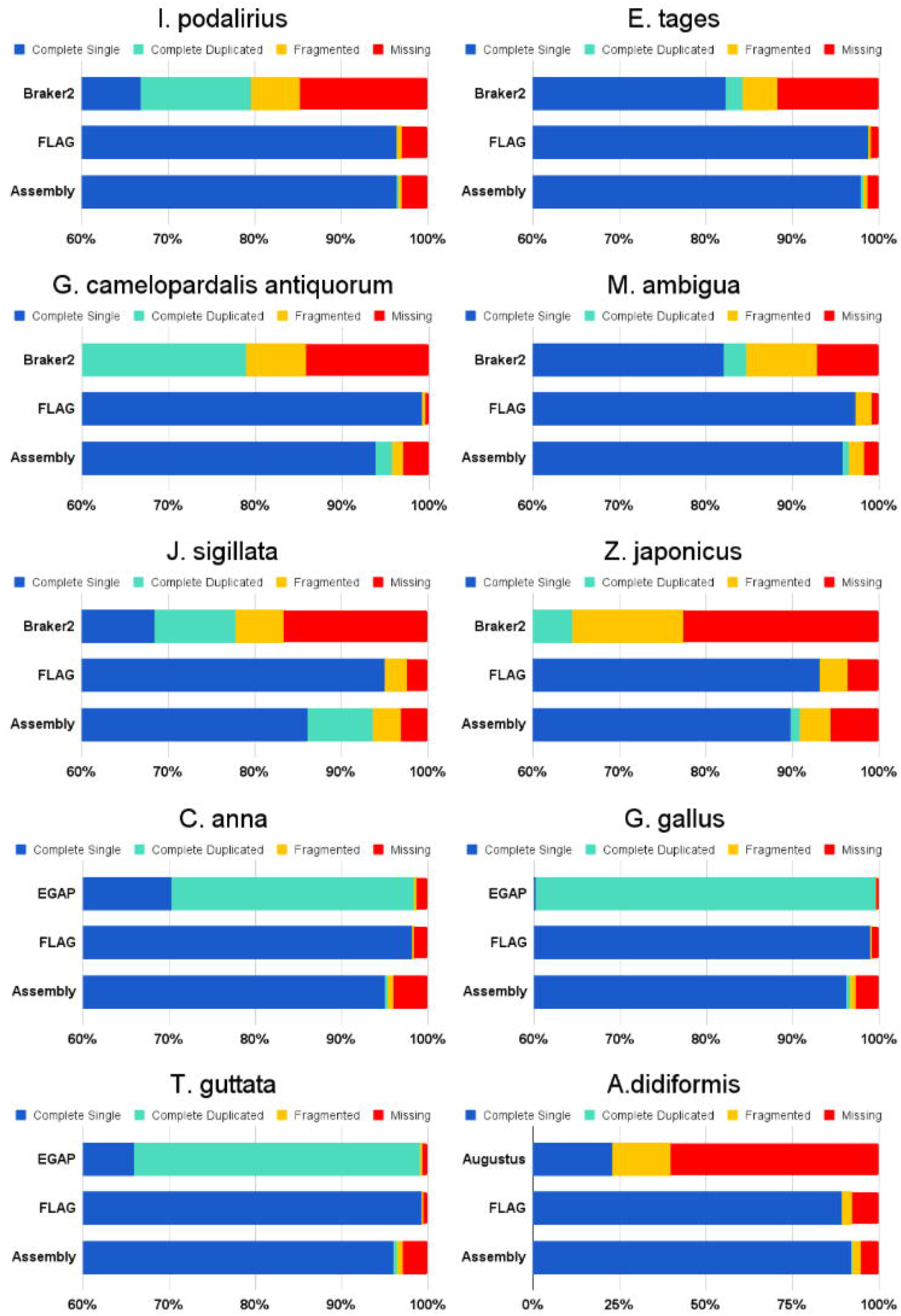
Overall workflow diagram of FLAG, not including the input genome as it is an input to all steps shown here. Some steps can be run in parallel and/or threaded to maximize CPU and RAM usage.

**Figure 2.**
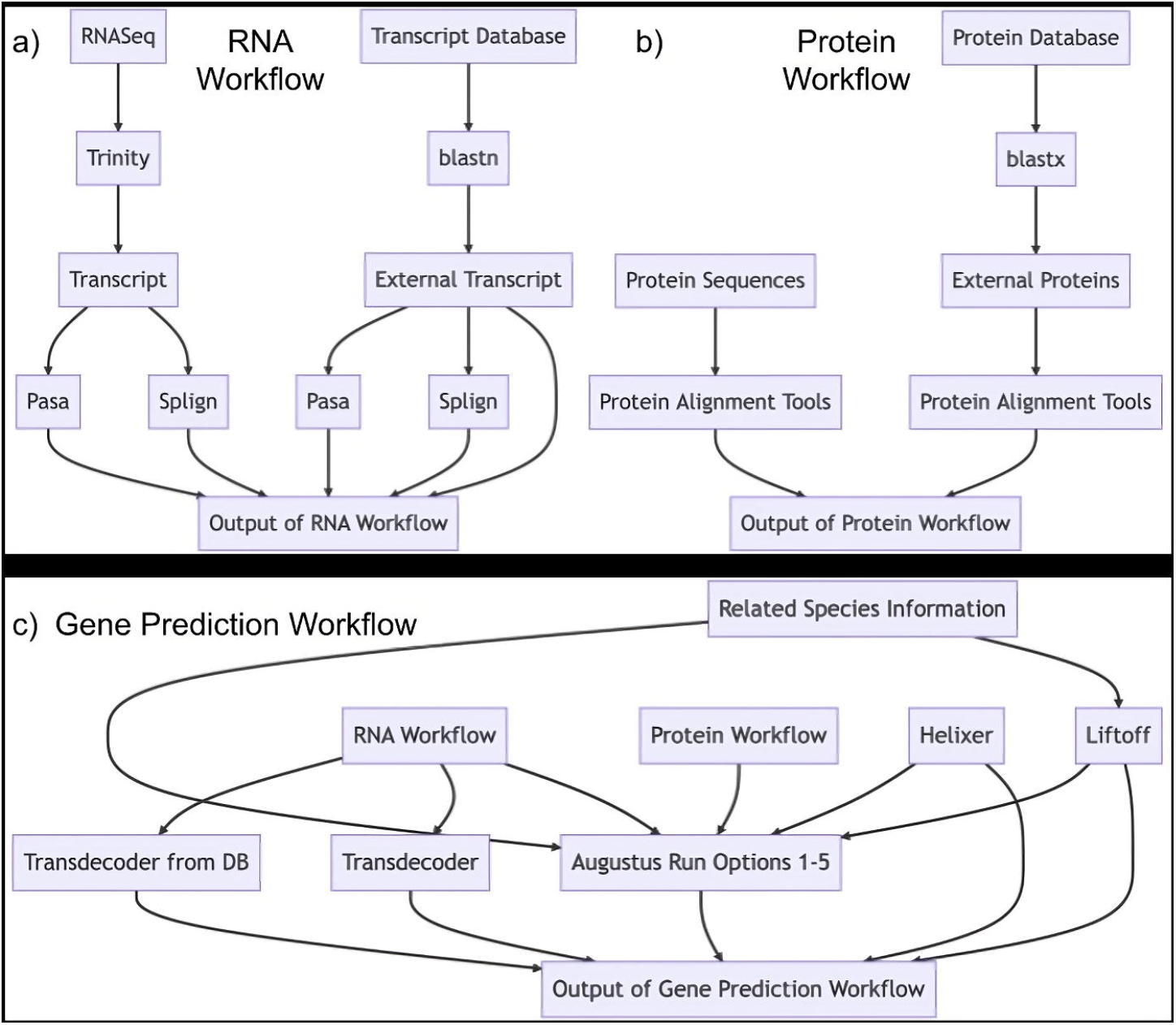
FLAG sub-workflows include a) the RNA, b) the protein, and c) the gene prediction workflows.

The RNA and protein workflows produce alignments to the target genome that can be used as evidence to train gene finders and as external evidence when determining gene confidence. The RNA workflow creates alignments of transcripts to the target genome assembly. RNASeq data is assembled into transcripts using Trinity (32). Assembled transcripts are aligned to the genome using Pasa (33) when from the same species or blast (34). Proteins are aligned to the genome using miniprot (35), genomethreader (36), exonerate (37), and/or prosplign (38).

The gene finding workflow produces the intron/exon structure of genes and includes the prediction of alternative transcripts. This workflow has four ab-initio gene finders implemented: Liftoff (17), Augustus (2), Transdecoder (39), and Helixer. Augustus can be trained with (i) RNA and protein alignments to the genomes (2), (ii) ab-initio gene predictions from Helixer or Liftoff. Otherwise, a pretrain model can be used from a closely related species. Transdecoder also uses RNA- and protein-to-genome alignments as evidence for ab-initio gene structural predictions. Helixer uses pre-trained models for vertebrates, invertebrates, land plants, and fungi. Models from these gene finders are combined using Evidence Modeler v2.0.0 (40). The gene structures from Evidence Modeler and all gene finders are filtered based on whether the prediction is a highly conserved gene using BUSCO and/or highly supported by evidence sources or multiple gene finders, with the heaviest weighting for evidence based support being on RNASeq/transcript data.

After completing the structural annotation, FLAG proceeds with the functional annotation of each predicted gene model using Eggnog-mapper (v2.1.12), with Eggnog (v5.0.2) or EnTAP integrated with Eggnog (v4.1). The outputs from both the structural and functional annotations are then merged into a comprehensive annotation, available to users in GTF, GFF3, and GBFF formats. Furthermore, FLAG employs BUSCO and AGAT to assess annotation quality. These tools examine gene completeness using orthologs, the total count of protein-coding genes, the prevalence of single-exon genes, and the average gene length.

### FLAG-Refiner

In order to maximize specificity of our gene predictions, we developed a new tool called FLAG-Refiner. FLAG-Refiner calculates the number of orthologs found in its database, how taxonomically close the taxids of these orthologs are to the target species, and the respective number of orthologs with exon counts similar to that of the gene annotation of interest. FLAG-Refiner has a large database constructed from every genome annotation in the RefSeq and EMBL databases, consisting of over 9,000 full genome annotations. For all of the genes in these annotations we calculate exon counts and organize genes, including isoforms if present, based on uniqueness, TaxaID, and assembly/annotation versions. Additionally, orthologs from the NCBI and EMBL databases are systematically cataloged, with the respective original ortholog database files being found at https://ftp.ncbi.nlm.nih.gov/gene/DATA/gene_orthologs.gz http://ftp.ensembl.org/pub/release-111/xml/ensembl-compara/homologies/Compara.111.protein_default.tree.orthoxml.xml.tar. When the database is being generated data from the NCBI and EMBL are first processed separately before being combined into a unified database. This unification step is done by relating genes from the NCBI RefSeq to those of EMBL via RefSeq to EMBL conversion tables, which can be found on the NCBI FTP site.

FLAG-Refine is then used to test gene predictions against several binary conditions, which can be modified, to determine the most likely true positive genes. Test 1 compares the exon counts of FLAG-predicted gene models with those of their corresponding orthologs, incorporating phylogenetic distances for accuracy. Test 2 calculates the percent exon coverage of each predicted gene by evidence alignments produced by Pasa, Splign, and miniprot. This percentage of what is deemed an acceptable alignment coverage for a gene is dependent on whether the gene is single- or multi-exon, the tool used to produce the alignment, and if the evidence is from an external database source or comes from evidence from a species that is closely related to the target species. As most of the over-annotation problems are related to over-prediction of single-exon genes, the default settings accept all multi-exon genes produced by FLAG. However, single-exon genes by default have requirements starting at 85% transcript-to-genome alignment exon coverage, a match to the orthoDB database, or at least 5 orthologs in the FLAG-Refiner database of which 20% must also be single-exon. From this, these settings can then iteratively relax themselves if the ratio of filtered single-to-multi-exon genes is under ratios deemed acceptable for the lineage in question or those set by the user. This relaxation consists of first allowing for the support of protein-to-genome alignments to deem genes as acceptable, starting at 100% exon coverage, before slowly decreasing the percentage of alignment support needed. This process is done until the settings that achieve the closest to acceptable single-to-multi-exon ratio are found. Excluded gene models are detailed in a separate report, providing justification for their removal and allowing for potential manual incorporation back into the final annotation. This transparency ensures that users understand the decision-making process and have the opportunity to adjust the annotation based on their expert judgment.

Metrics produced by FLAG-Refiner added to the final annotation include the number of orthologs identified, the distribution of single versus multi-exon genes, alignment coverage percentages, and congruence with orthoDB data. These enriched final annotations, available in GTF, GFF3, and GBFF formats, undergo a final round of quality assessment with BUSCO and AGAT, culminating in a comprehensive annotation report. This report highlights the improvements made by incorporating FLAG-Refiner and assures users of the robustness and reliability of the genome annotation.

## Results

In our comprehensive evaluation of the newly developed “Find, Label, Annotate Genomes” (FLAG) annotation pipeline, we systematically assessed its performance across a spectrum of genomic complexities and data availability scenarios to demonstrate its versatility and robustness. Our analyses encompass four distinct tests. First, we benchmarked the accuracy and computational efficiency of different tools used in FLAG to determine the default tools, focusing on two with particularly noteworthy outcomes. Second, we examined FLAG’s capability to annotate non-model organism genomes without relying on species-specific evidence, highlighting its utility in genomic research where such data is scarce or unavailable. Third, we validated FLAG’s performance on model organism genomes for which species-specific evidence was accessible, ensuring the tool’s effectiveness in more controlled research environments.

Lastly, we presented a challenging test case involving the highly fragmented genome of the extinct Moa bird, *A. didiformis*, which lacks any species-specific evidence. This series of tests underscores FLAG’s potential to significantly advance genomic annotation across diverse research contexts.

### Non-model Highly-fragmented Genome Assemblies

Using FLAG, we annotated 6 non-model species that also had published BRAKER2 annotations: *E. tages* (Dingy skipper Butterfly), *I. podalirius* (Scarce Swallowtail Butterfly), *J. sigillata* (Iron Walnut Tree), *M. ambigua* (Golden Perch), *Z. japonicus* (Japanese white-eye), and *G. camelopardalis antiquorum* (Kordofan Giraffe). For these species, all FLAG runs used no species-specific evidence regardless of whether the BRAKER2 annotations did.

Most of these model genomes are highly fragmented with an average of 28,093 contigs (Table 1). Compared to BRAKER2, for these genome assemblies, FLAG annotations have on average 18% higher complete BUSCO scores, 6.68% fewer fragmented genes, 12.68% fewer missing genes and 17x lower error rate in protein-coding predicted gene counts (Table 2).

**Table 1.** Assembly statistics on the genomes used for the FLAG annotation tests.

| Genome | Size (Mb) | Scaffolds | Scaffold N50 | Scaffold L50 | Contigs | Contig N50 | Contig L50 |
| --- | --- | --- | --- | --- | --- | --- | --- |
| *E. tages* | 329 | 41 | 11,903,105 | 13 | 111 | 8,006,790 | 16 |
| *I. podalirius* | 430 | 260 | 15,090,856 | 13 | 446 | 5,183,580 | 26 |
| *J. sigillata* | 585 | 6,413 | 238,362 | 699 | 54,794 | 46,167 | 3,256 |
| *M. ambigua* | 661 | 7,165 | 252,501 | 730 | 7,352 | 243,852 | 757 |
| *Z. japonicus* | 1,005 | 29,442 | 9,685,185 | 29 | 73,843 | 38,076 | 7,569 |
| *G. camelopardalis* | 2,963 | 10,445 | 27,504,022 | 25 | 32,009 | 228,443 | 3,139 |
| *antiquorum* |  |  |  |  |  |  |  |
| *C. anna* | 1,073 | 159 | 74,081,004 | 4 | 584 | 14,522,327 | 21 |
| *G. gallus* | 1,067 | 213 | 90,861,225 | 4 | 676 | 18,834,961 | 18 |
| *T. guttata* | 1,070 | 198 | 70,982,421 | 6 | 550 | 8,964,551 | 32 |
| *A. didiformis* | 1,190 | 1,909 | 3,350,051 | 111 | 1,734,944 | 823 | 321,557 |

**Table 2.** Gene annotation statistics of BRAKER2 and FLAG against respective NCBI comparison species being what is deemed as expected truth, shaded in gray. Statistics were calculated using AGAT v1.0.0. For *A. didiformis* the closest related species was *T. guttatus*, however, the assembly for *A. didiformis* was built off of *D. novaehollandiae* and as such the current *A. didiformis* assembly should resemble a mix between the two.

| Genome | Annotation Method | # Protein Coding Genes | # Single Exon Genes | Mean Gene Length |
| --- | --- | --- | --- | --- |
| *E. tages* | BRAKER2 | 15,237 | 2,618 | 6,721 |
| *E. tages* | FLAG | 12,929 | 1,784 | 10,948 |
| *E. tages* | FLAG w/ Refiner | 12,283 | 1,138 | 11,471 |
| *I. podalirius* | BRAKER2 | 17,826 | - | - |
| *I. podalirius* | FLAG | 12,477 | 1,669 | 12,092 |
| *I. podalirius* | FLAG w/ Refiner | 11,830 | 1,022 | 12,722 |
| *D. plexippus*<br>(*E. tages* and *I. podalirius* comparison) | NCBI | 13,105 | 1,015 | 10,501 |
| *G. camelopardalis antiquorum* | BRAKER2 | 59,082 | 25,710 | 9,869 |
| *G. camelopardalis antiquorum* | FLAG | 20,532 | 2,258 | 45,055 |
| *G. camelopardalis antiquorum* | FLAG w/ Refiner | 20,532 | 2,258 | 45,055 |
| *C. elaphus* (G. | NCBI | 22,941 | 3,251 | 49,621 |
| *camelopardalis antiquorum* comparison) |  |  |  |  |
| *M. ambigua* | BRAKER2 | 36,108 | 5,000 | 6,296 |
| *M. ambigua* | FLAG | 25,367 | 2,172 | 12,216 |
| *M. ambigua* | FLAG w/ Refiner | 23,350 | 1,155 | 12,706 |
| *M. salmoides* (*M. ambigua* comparison) | NCBI | 27,179 | 1,088 | 19,634 |
| *Z. japonicus* | BRAKER2 | 71,966 | 6,090 | 4,120 |
| *Z. japonicus* | FLAG | 18,012 | 2,807 | 23,102 |
| *Z. japonicus* | FLAG w/ Refiner | 16,013 | 808 | 25,936 |
| *H. rustica* (*Z. japonicus* comparison) | NCBI | 15,516 | 580 | 35,255 |
| *J. sigillata* | BRAKER2 | 89,512 | 10,653 | 2,993 |
| *J. sigillata* | FLAG | 34,806 | 8,982 | 4,026 |
| *J. sigillata* | FLAG w/ Refiner | 31,833 | 6,009 | 4,359 |
| *J. microcarpa* x *J. regia* (*J. sigillata* comparison) | NCBI | 27,962 | 4,238 | 5,586 |
| *J. regia* (*J. sigillata* comparison) | NCBI | 30,709 | 5,255 | 5,309 |
| *C. anna* | FLAG | 15,377 | 972 | 30,591 |
| *C. anna* | FLAG w/ Refiner | 15,064 | 890 | 31,073 |
| *C. anna* | NCBI | 14,711 | 564 | 34,495 |
| *G. gallus* | FLAG | 18,996 | 1,665 | 26,036 |
| *G. gallus* | FLAG w/ Refiner | 18,864 | 1,533 | 26,216 |
| *G. gallus* | NCBI | 18,023 | 806 | 33,309 |
| *T. guttata* | FLAG | 17,171 | 1,219 | 28,226 |
| *T. guttata* | FLAG w/ Refiner | 16,942 | 1,127 | 28,512 |
| *T. guttata* | NCBI | 16,520 | 587 | 33,996 |
| *A. didiformis* | Augustus (3 runs) | 9,797-13,763 | 106-122 | 24,369-35,722 |
| *A. didiformis* | FLAG | 14,329 | 756 | 30,972 |
| *A. didiformis* | FLAG w/ Refiner | 14,329 | 756 | 30,972 |
| *D. novaehollandiae* (*A. didiformis* comparison) | NCBI | 15,554 | 552 | 37,263 |
| *T. guttatus* (*A. didiformis* comparison) | NCBI | 15,723 | 820 | 22,119 |

### Model Avian Genome Assemblies

We used FLAG to predict gene annotations in 3 avian model organisms, using species-specific evidence, and compared our annotation to the publicly available NCBI annotation (Table 1). These model species were *C. anna* (Anna’s Hummingbird), *G. gallus* (Chicken), and *T. guttata* (Zebra Finch).

Resulting gene predictions from FLAG were concordant with NCBI annotations using the number of single- and multi-exon genes, mean gene lengths, and BUSCO scores. EGAP predicted an average 16,418 protein-coding genes and 98.67% complete gene orthologs found, using the Aves ODB10 database. FLAG with FLAG-Refiner predicted an average 16,957 protein-coding genes and 98.93% complete Aves ODB10 BUSCOs (Figure 3 and Table 2).The main difference between these annotations is that FLAG produces on average 531 more single exon genes when compared to EGAP. All of these extra genes were supported by FLAG-Refiner and obeyed at least one of the following requirements: (i) 90% exon coverage by transcript to genome alignments, (ii) 100% exon coverage by protein to genome alignments, (iii) a unique orthoDB match, and/or (iv) at least 5 single-exon orthologs. Out of all of the extra single exon genes predicted by FLAG, for these three model species, 14% met at least one of these requirements, 54% met two, 26% met three, and 6% met all four requirements.

**Figure 3.**
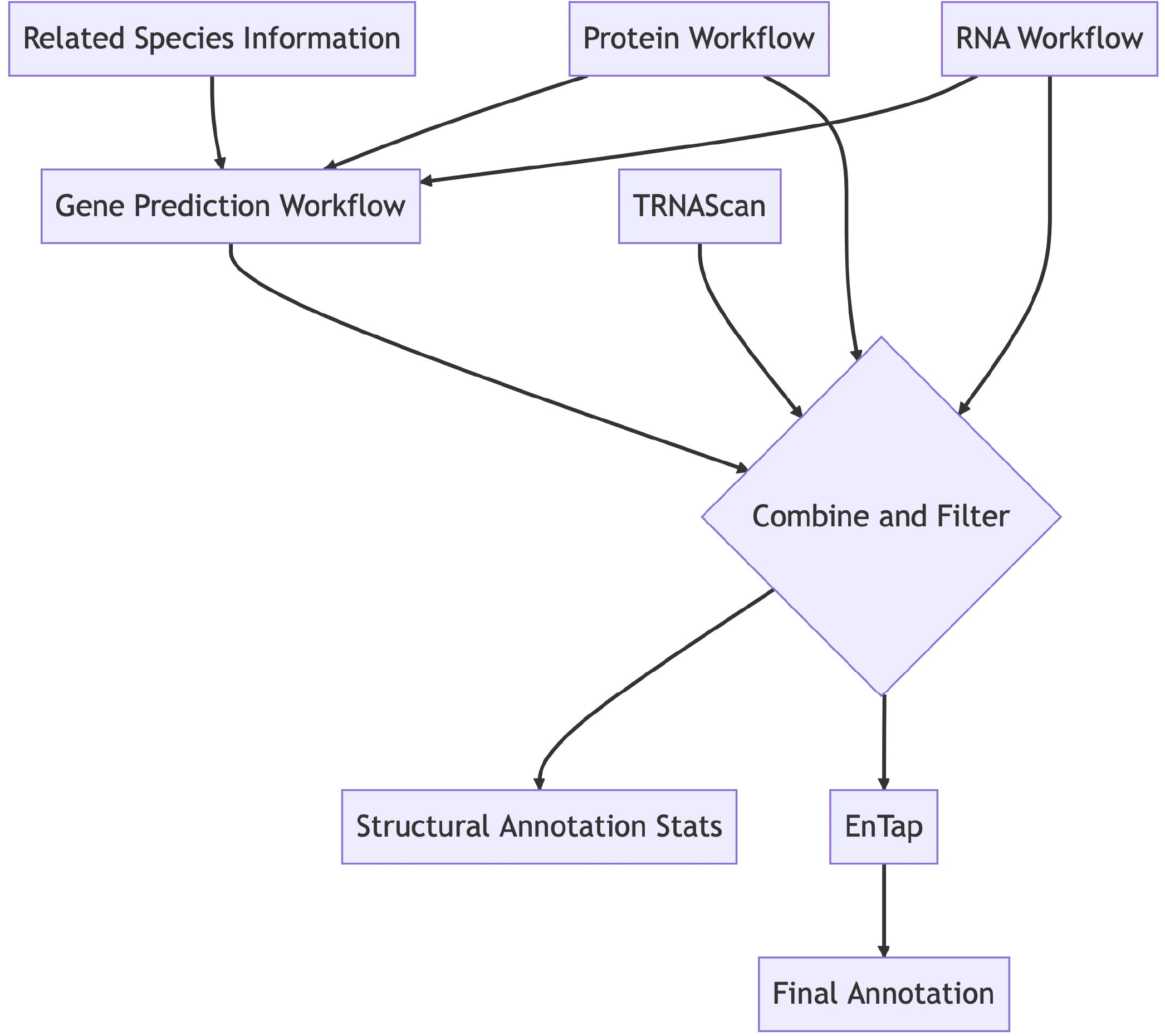
BUSCO scores of BRAKER2, Augustus, and FLAG annotations compared against their respective genome assembly BUSCO scores. Lineages used include Lepidoptera for *I. podalirius* and *E. tages*, Eudicots for *J. sigillata*, Actinopterygii for *M. ambigua*, Sauropsida for *Z. japonicus and A. didiformis*, Mammalia for *G. camelopardalis antiquorum, and Aves for C. anna, G. gallus*, and *T. guttata*. All BUSCO scores were calculated with BUSCO v5.3.2 and ortho DB v10 (44).

### Extinct Moa Bird Genome Assembly

We used FLAG to annotate a highly fragmented genome from an extinct species, *Anomalopteryx didiformis* (Little-bush Moa) (28), whose genome assembly has over 1.7 million contigs (Table 1). This specimen lacks RNA and protein data and was assembled from degraded DNA. Because there is no publicly available genome, we compared our annotation to that of its closest sequenced and annotated relatives *D. novaehollandiae* (Emu) and *T. guttatus* (Zebra Finch; Figure 3 and Table 2). We also annotated this genome with AUGUSTUS alone. Compared to AUGUSTUS, FLAG predicted 66.3% more complete BUSCO genes, 13.8% less fragmented genes, and 52.5% fewer missing genes (Figure 3). The number of protein-coding genes predicted by FLAG was similar to those annotated in the closely related species (Table 2).

### Benchmarking of Tools in FLAG Defaults

During the evaluation with non-model genomes the robustness of FLAG was highly tested as some of the closely related species evidence was not similar enough to the species of interest to provide high-quality alignments, as can be the case with non-model genomes. For instance, in butterflies, when using the *D. plexippus* genome as input evidence in Liftoff to create annotations for *I. podalirius* and *E. tages* genomes, Liftoff only predicted 5,000 genes of ∼13k expected for each species, since *D. plexippus* mapped quite poorly to the genome assemblies being annotated. However, we found that even when one of the main gene prediction tools, in this case, Liftoff, produces results drastically different from the expected, the final annotation remains relatively unaffected. In the same example, the total protein-coding gene count of the final FLAG results with and without Liftoff only differed by a maximum of 21 genes (Supplementary Table 2).

Augustus was also tested with FLAG, which was run with all of the normal defaults, including and excluding Augustus, for our 6 non-model species and the Moa. This resulted in slightly more variability in the total number of predicted protein-coding genes when compared to the Liftoff test. The average difference in the number of predicted protein-coding genes for FLAG runs with and without Augustus was 168 for the 6 non-model species, and 1562 genes for the Moa (Supplementary Table 3). This difference is likely highly related to the quality of the target genome assembly, so our recommendation is for Augustus always to be run for low-quality assemblies and optional for higher-quality genome assemblies. While Augustus runs are not deemed as hindering results in any cases for some it may be beneficial to leave it out as this can decrease run time anywhere from a few hours up to a whole day, for large genomes.

Typical run times using FLAG default run parameters with proteins, transcripts, and a related species annotation as evidence depend on data size. For genomes between 800 Mb and 3.3GB typical FLAG runs can take from 3 hours to 4 days. This can increase relatively greatly if unassembled RNASeq evidence is input, however, run time will largely depend on the amount of RNASeq evidence with a reported run time of 4-9 minutes per million pair-reads (41). Though to speed this up FLAG can run trinity in parallel on separate tissue or specimen samples, as is recommended if over 200 GB of raw RNASeq evidence is provided.

This fast speed is partially due to the use of miniprot as the primary tool for the traditionally computationally expensive protein-to-genome alignment step. This decision was based on our findings when aligning *D. melanogaster* proteins obtained from the NCBI to the *D. melanogaster* genome. Therein, we found that miniprot significantly reduced protein-to-genome alignment time by up to 430 times compared to exonerate and 655 times compared to genomethreader, all while maintaining similar levels of accuracy. In contrast, on the gene prediction side, Transdecoder demonstrated quick processing times in our assessments, but it lagged in accuracy, emerging as the least effective among the gene predictors we implemented.

## Discussion

In this work, we introduce the Find, Label, Annotate Genomes (FLAG) pipeline with FLAG-Refiner, a cutting-edge toolkit designed for the comprehensive annotation of genomes across a diverse array of species. FLAG distinguishes itself through the integration of advanced gene-finding algorithms such as Helixer and Liftoff along with detailed gene model filtering, both internal to FLAG and FLAG-Refiner, ensuring both swift and precise annotations. A standout feature of FLAG is its versatility: available as open-source software for those preferring hands-on customization as well as along a user-friendly, point-and-click interface, complete with an array of pre-curated databases, through the Form Bio platform for ease of use and efficiency.

One of the core strengths of FLAG lies not just in the accuracy of its structural annotations but also in its all-encompassing approach where it also produces functional annotations and provides annotation files in multiple formats that contain both annotation types together. Along with this it also generates detailed reports using BUSCO and AGAT, to generate annotation statistics analogous to GenomeQC (42), and FLAG-Refiner so that every annotation can be examined in detail. This approach provides users with a nuanced assessment of gene prediction quality, advancing the rigorous standards of genome assembly and annotation quality control.

FLAG’s effectiveness and utility are further underscored by its high-quality results when annotating both low- and high-quality input, allowing for its application to over 70 avian species genomes in just the past six months alone (43). This achievement demonstrates the pipeline’s scalability and reliability in handling genomic data from a broad spectrum of organisms, from the well-studied to the rare and extinct. Notably, FLAG’s proficiency was tested with the highly fragmented genome of the extinct Moa bird, *A. didiformis*, for which it successfully generated high-quality annotations from ancient DNA in the absence of species-specific evidence. This feat highlights FLAG’s capability to navigate the challenges presented by incomplete or highly fragmented genomes, offering valuable insights into the genetic makeup of both living and lost species.

In conclusion, FLAG and FLAG-Refiner represent a significant leap forward in genomic research tools, combining speed, accuracy, flexibility, and user accessibility. Their proven track record, underscored by their application to a wide range of species, positions FLAG as an ideal resource for geneticists, evolutionary biologists, and conservation scientists alike, facilitating a deeper understanding of the genomic intricacies of diverse biological lineages in the age of large-scale cross-species genomics.

### Software Availability Statement

FLAG is available and open source at https://github.com/formbio/FLAG

