## Supplementary Table 1, Supplementary Figure 1, Supplementary Figure 2 for "Find, Label, Annotate Genomes: FLAG, a fully automated tool for structural and functional gene annotation"

**Supplementary Figures:**

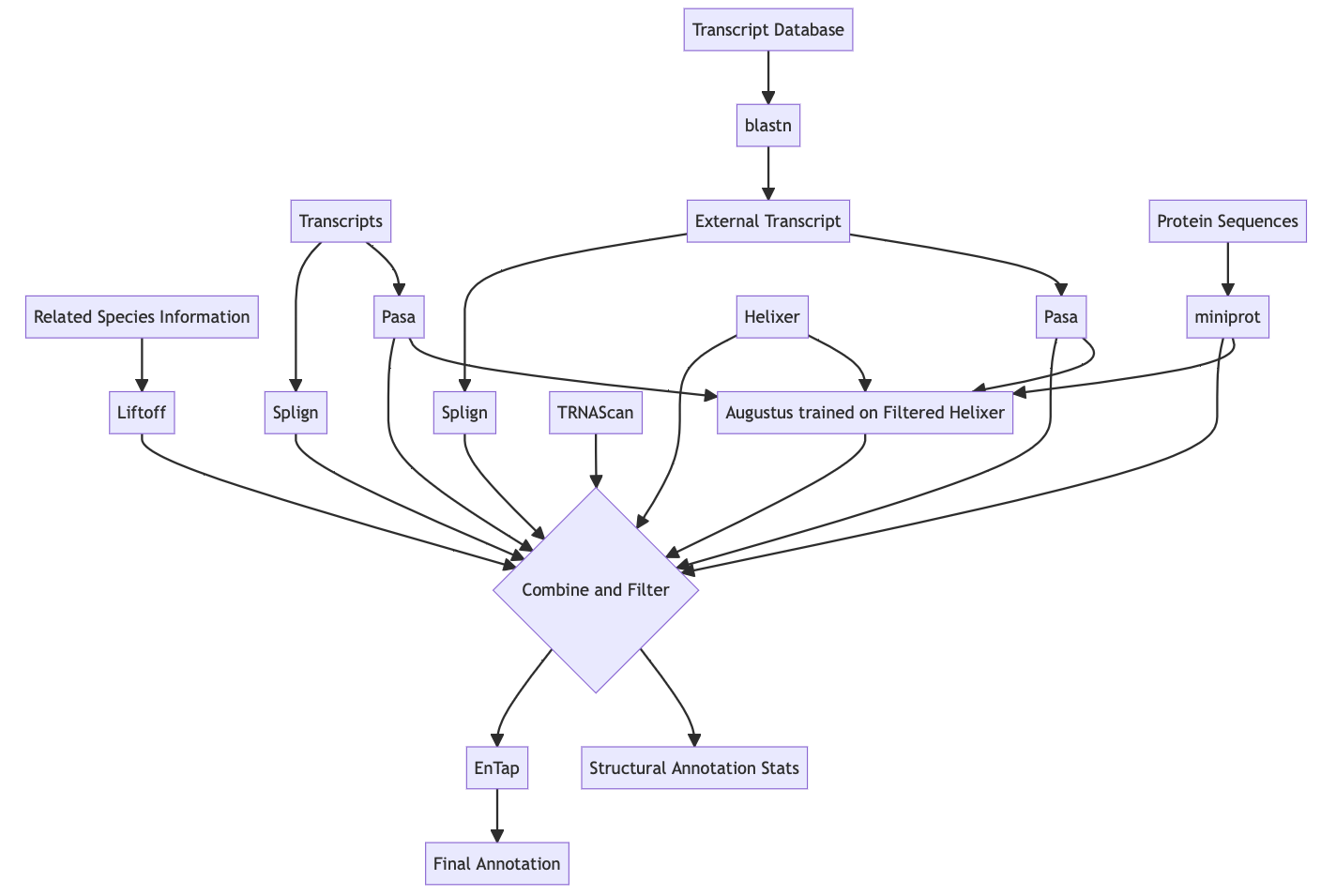

Supplementary Figure 1. Default run configuration of FLAG

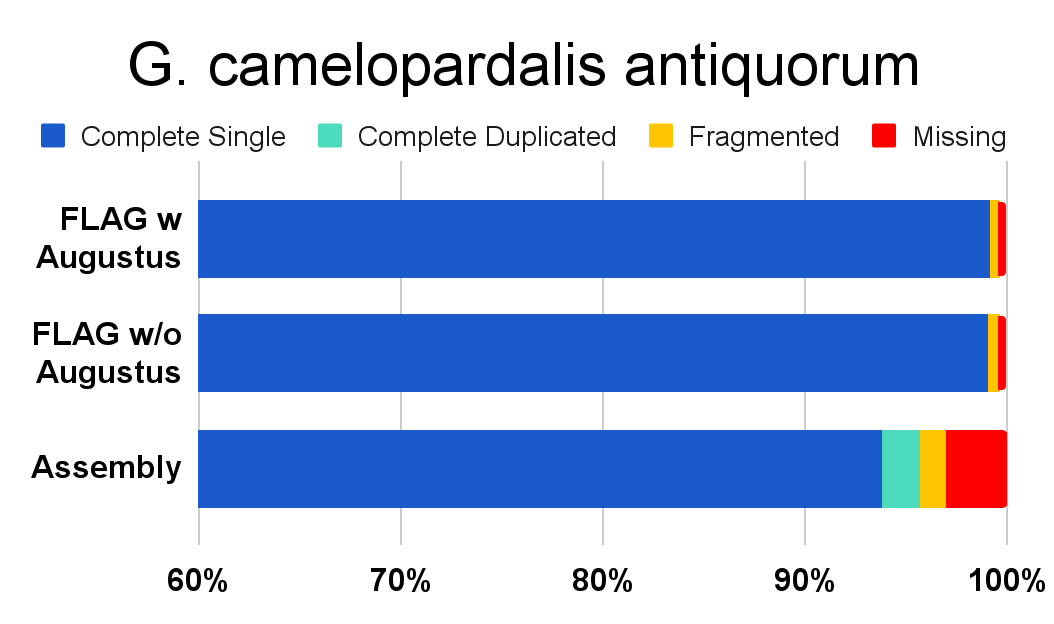

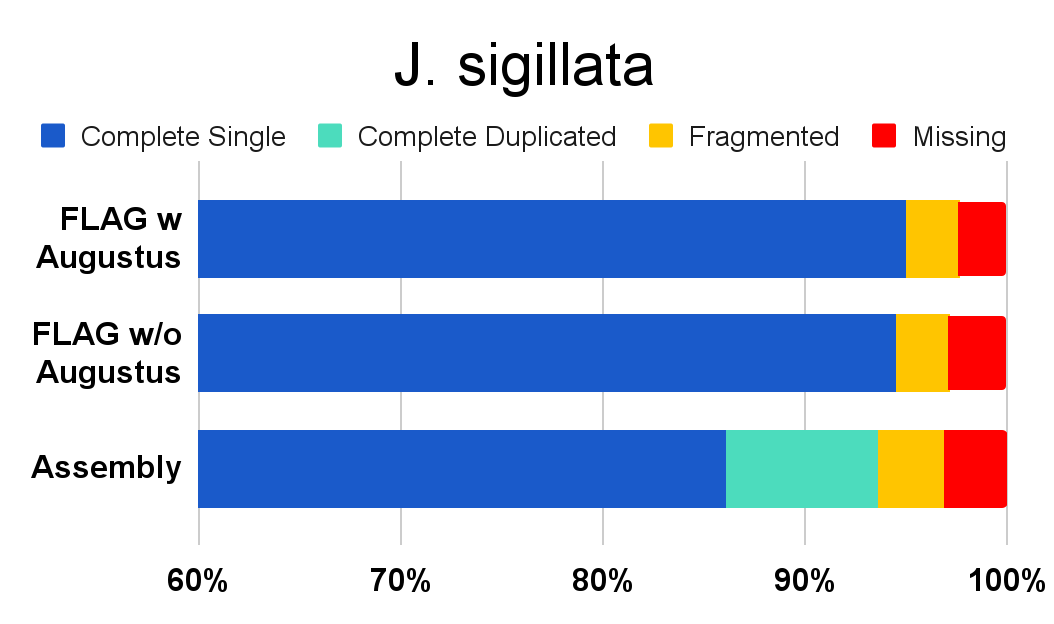

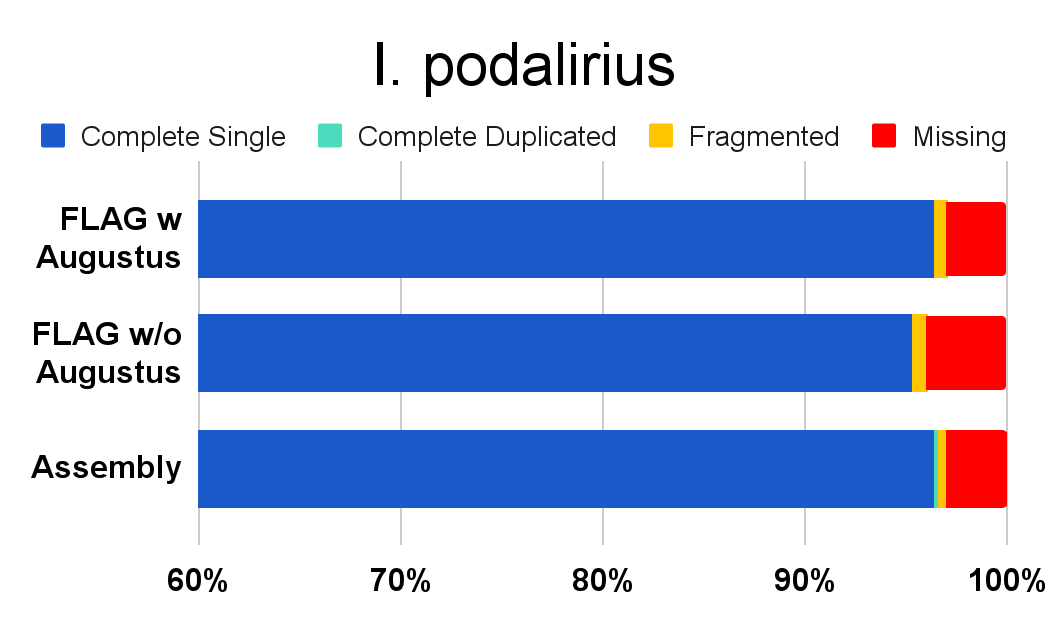

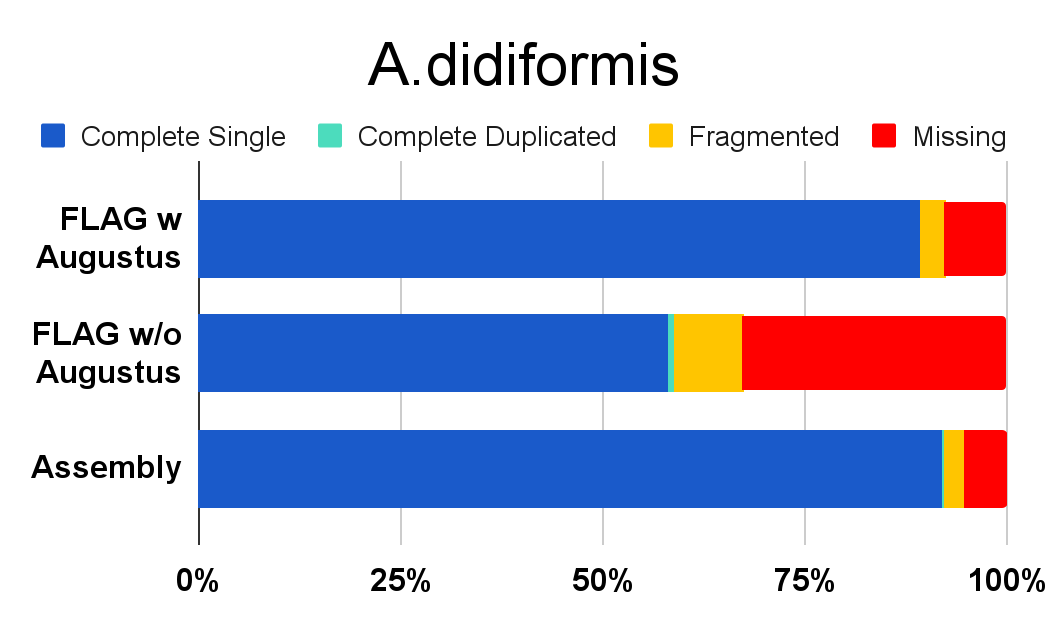

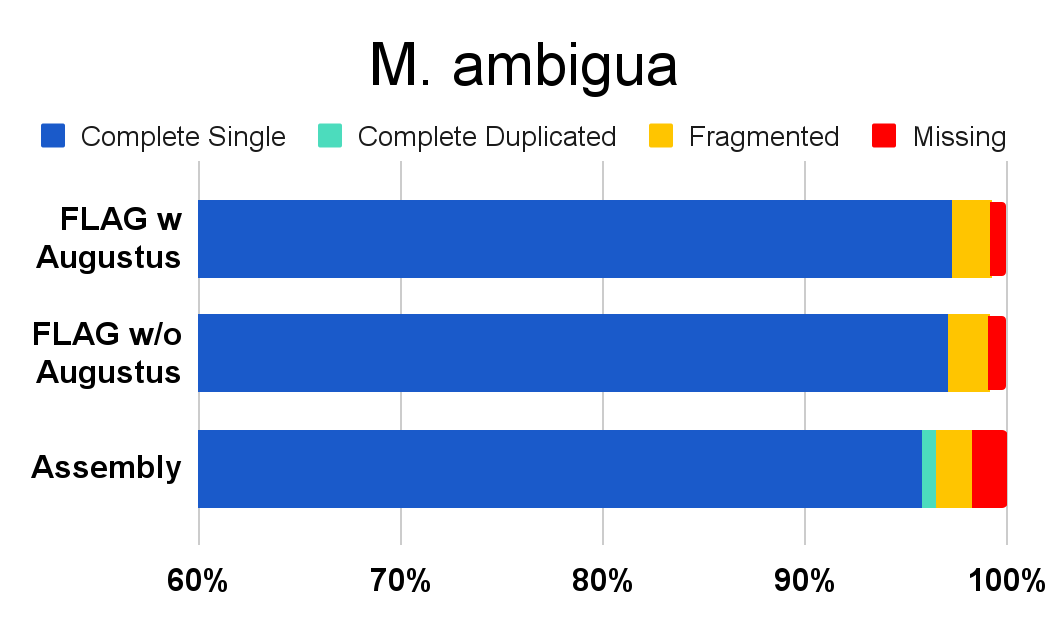

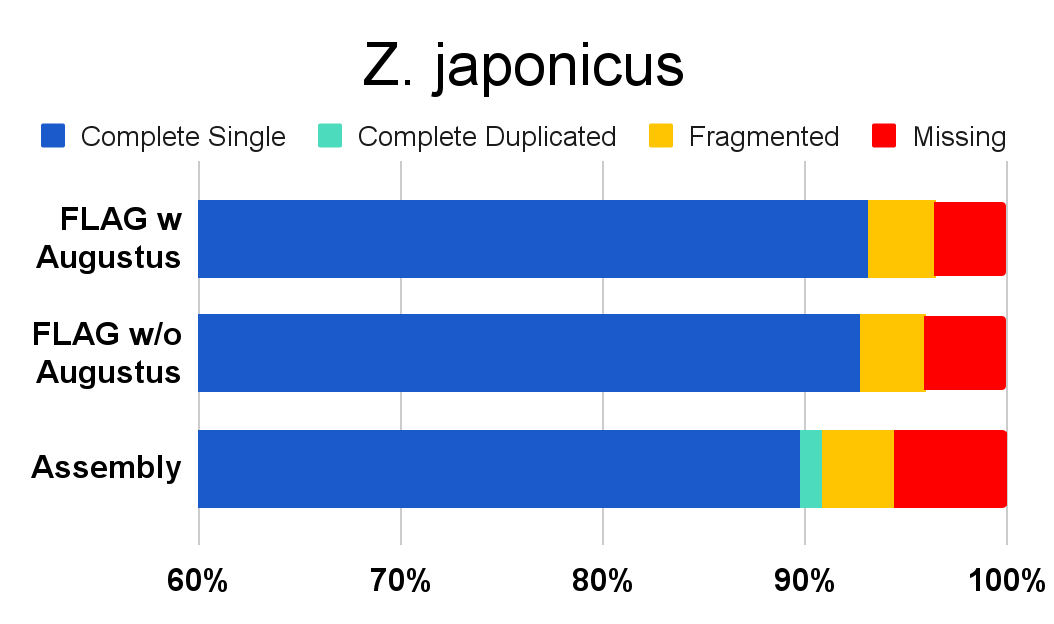

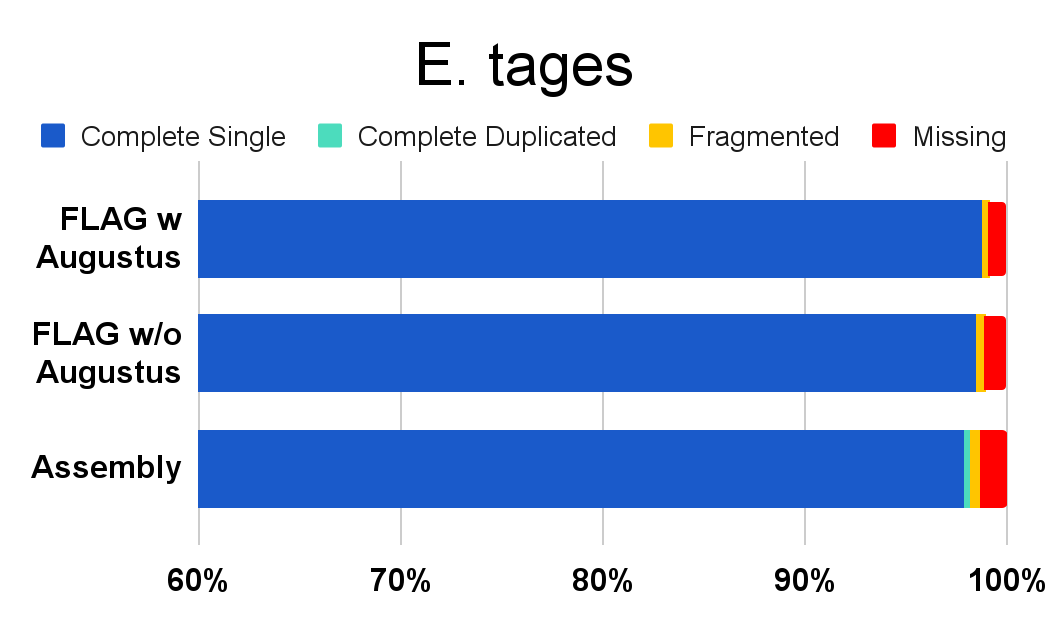

Supplementary Figure 2. BUSCO scores of FLAG with and without Augustus annotations compared against their respective genome BUSCO scores. Lineages used include lepidoptera odb10 for *I. podalirius* and *E. tages*, eudicots odb10 for *J. sigillata*, actinopterygii odb10 for *M. ambigua*, and sauropsida odb10 for *Z. japonicus* and *A. didiformis*, and mammalia odb10 for *G. camelopardalis antiquorum*. All BUSCO scores were calculated with BUSCO v5.3.2 and ortho DB v10.

**Supplementary Tables:**

Supplementary Table 1. NCBI RefSeq accession numbers for genomes and annotations used within the study for annotation references, tool speed tests, and as evidence for annotations.

| **Species** | **ResSeq Accession** | **Evidence for Species** | **Comparison for Species** |
| --- | --- | --- | --- |
| *Drosophila melanogaster* | GCF_000001215.4 | - | - |
| *Danaus plexippus plexippus* | GCF_009731565.1 | *E. tages and I. podalirius* | *E. tages and I. podalirius* |
| *Leguminivora glycinivorella* | GCF_023078275.1 | *E. tages and I. podalirius* | - |
| *Pieris napi* | GCF_905475465.1 | *E. tages and I. podalirius* | - |
| *Zerene cesonia* | GCF_012273895.1 | *E. tages and I. podalirius* | - |
| *Bos taurus* | GCF_000003205.7 | *G. camelopardalis antiquorum* | - |
| *Bubalus bubalis* | GCF_019923935.1 | *G. camelopardalis antiquorum* | - |
| *Budorcas taxicolor* | GCF_023091745.1 | *G. camelopardalis antiquorum* | - |
| *Cervus canadensis* | GCF_019320065.1 | *G. camelopardalis antiquorum* | - |
| *Cervus elaphus* | GCF_910594005.1 | *G. camelopardalis antiquorum* | *G. camelopardalis antiquorum* |
| *Ovis aries* | GCF_002742125.1 | *G. camelopardalis antiquorum* | - |
| *Micropterus dolomieu* | GCF_021292245.1 | *M. ambigua* | - |
| *Micropterus salmoides* | GCF_014851395.1 | *M. ambigua* | *M. ambigua* |
| *Siniperca chuatsi* | GCF_020085105.1 | *M. ambigua* | - |
| *Hirundo rustica* | GCF_015227805.2 | *Z. japonicus* | *Z. japonicus* |
| *Manacus vitellinus* | GCF_001715985.3 | *Z. japonicus* | - |
| *Passer montanus* | GCF_014805655.1 | *Z. japonicus* | - |
| *Pipra filicauda* | GCF_003945595.2 | *Z. japonicus* | - |
| *Juglans microcarpa x Juglans regia* | GCF_004785595.1 | *J. sigillata* | *J. sigillata* |
| *Juglans regia* | GCF_001411555.2 | *J. sigillata* | *J. sigillata* |
| *Populus trichocarpa* | GCF_000002775.5 | *J. sigillata* | - |
| *Quercus robur* | GCF_932294415.1 | *J. sigillata* | - |
| *Gallus gallus* | GCF_016699485.2 | *A. didiformis and G. gallus* | *G. gallus* |
| *Struthio camelus* | GCF_000698965.1 | *A. didiformis* | - |
| *Tinamus guttatus* | GCF_003957565.2 | *A. didiformis and T. guttatus* | *A. didiformis and T. guttatus* |
| *Dromaius novaehollandiae* | GCF_003342905.1 | *A. didiformis* | *A. didiformis* |
| *Calypte anna* | GCF_003957555.1 | *C. anna* | *C. anna* |

Supplementary Table 2. Annotation statistics of *I. podalirius* and *E. tages* of FLAG run both with and without Liftoff.

| **Genome** | **Annotation Method** | **# Protein Coding Genes** | **# Single Exon Genes** | **Mean Gene Length** |
| --- | --- | --- | --- | --- |
| *E. tages* | FLAG with Liftoff | 12,929 | 1,784 | 10,948 |
| *E. tages* | FLAG without Liftoff | 12,908 | 1,765 | 10,951 |
| *I. podalirius* | FLAG with Liftoff | 12,466 | 1,671 | 11,979 |
| *I. podalirius* | FLAG without Liftoff | 12,461 | 1,658 | 11,988 |

Supplementary Table 3. The total number of protein-coding genes, single exon protein-coding genes, and mean gene length of the full FLAG defaults both with and without Augustus.

| **Genome** | **Annotation Method** | **# Protein Coding Genes** | **# Single Exon Genes** | **Mean Gene Length** |
| --- | --- | --- | --- | --- |
| *E. tages* | FLAG w Augustus | 12,929 | 1,784 | 10,948 |
| *E. tages* | FLAG w/o Augustus | 13,034 | 1,531 | 11,187 |
| *I. podalirius* | FLAG w Augustus | 12,477 | 1,669 | 12,092 |
| *I. podalirius* | FLAG w/o Augustus | 12,569 | 1,395 | 12,324 |
| *D. plexippus*  (*E. tages* and *I. podalirius* comparison) | NCBI | 13,105 | 1,015 | 10,501 |
| *G. camelopardalis antiquorum* | FLAG w Augustus | 20,532 | 2,258 | 45,055 |
| *G. camelopardalis antiquorum* | FLAG w/o Augustus | 20,539 | 2,336 | 44,574 |
| *C. elaphus* (*G. camelopardalis antiquorum* comparison) | NCBI | 22,941 | 3,251 | 49,621 |
| *M. ambigua* | FLAG w Augustus | 25,367 | 2,172 | 12,216 |
| *M. ambigua* | FLAG w/o Augustus | 25,439 | 1,677 | 13,007 |
| *M. salmoides* (*M. ambigua* comparison) | NCBI | 27,179 | 1,088 | 19,634 |
| *Z. japonicus* | FLAG w Augustus | 18,012 | 2,807 | 23,102 |
| *Z. japonicus* | FLAG w/o Augustus | 18,224 | 1,537 | 23,056 |
| *H. rustica* (*Z. japonicus* comparison) | NCBI | 15,516 | 580 | 35,255 |
| *J. sigillata* | FLAG w Augustus | 34,806 | 8,982 | 4,026 |
| *J. sigillata* | FLAG w/o Augustus | 35,326 | 8,581 | 3,912 |
| *J. microcarpa x J. regia* (*J. sigillata* comparison) | NCBI | 27,962 | 4,238 | 5,586 |
| *J. regia* (*J. sigillata* comparison) | NCBI | 30,709 | 5,255 | 5,309 |
| *A. didiformis* | FLAG w Augustus | 14,329 | 756 | 30,972 |
| *A. didiformis* | FLAG w/o Augustus | 12,767 | 945 | 28,971 |
| *D. novaehollandiae* (*A. didiformis* comparison) | NCBI | 15,554 | 552 | 37,263 |
| *T. guttatus* (*A. didiformis* comparison) | NCBI | 15,723 | 820 | 22,119 |
